# Phosphoinositides regulate the TRPV1 channel via two functionally distinct binding sites

**DOI:** 10.1101/558874

**Authors:** Aysenur Torun Yazici, Eleonora Gianti, Marina A. Kasimova, Vincenzo Carnevale, Tibor Rohacs

## Abstract

Regulation of the heat- and capsaicin-activated Transient Receptor Potential Vanilloid 1 (TRPV1) channel by phosphoinositides is controversial. In a recent cryoEM structure, an endogenous phosphoinositide was detected in the vanilloid binding site, and phosphoinositides were proposed to act as competitive vanilloid antagonists. This model is difficult to reconcile with phosphatidylinositol 4,5- bisphosphate [PtdIns(4,5)P_2_] being a well established positive regulator of TRPV1. To resolve this controversy, we propose that phosphoinositides regulate TRPV1 via two functionally distinct binding sites. Our molecular dynamics simulations show that phosphatidylinositol (PtdIns) is more stable in the vanilloid binding site, whereas a distinct site responsible for activation is preferentially occupied by PtdIns(4,5)P_2_. Consistently, we show that in the presence of PtdIns(4,5)P_2_ in excised patches PtdIns partially inhibited TRPV1 activity induced by low, but not high capsaicin concentrations. In the absence of PtdIns(4,5)P_2_ on the other hand, PtdIns partially stimulated TRPV1 activity presumably by binding to the activating site. Overall, our data resolve a major controversy in the regulation of TRPV1.

## INTRODUCTION

Transient Receptor Potential Vanilloid 1 (TRPV1) is a heat- and capsaicin-activated Ca^2+^ permeable non-selective cation channel expressed in primary sensory neurons of the Dorsal Root Ganglia (DRG) and Trigeminal Ganglia (TG) (Caterina et al., 1999). Stimulation of cell surface receptors by pro-inflammatory mediators sensitizes this channel to heat and capsaicin, a phenomenon thought to underlie the increased sensitivity to heat upon inflammation. Accordingly, genetic deletion of this channel in mice essentially eliminated thermal hyperalgesia (Caterina et al., 2000).

Phosphoinositides, especially PtdIns(4,5)P_2_, are general regulators of many ion channels, including those belonging to the transient receptor potential (TRP) family (Rohacs, 2014). In most cases, PtdIns(4,5)P_2_ acts as a necessary cofactor for channel activity (Suh and Hille, 2008). Phosphoinositides modulate TRPV1 activity in a complex manner; both positive and negative effects of these lipids have been proposed (Rohacs, 2015). It was firstly suggested that phosphatidylinositol 4,5-bisphosphate [PtdIns(4,5)P_2_] negatively regulates TRPV1, and that relief from this inhibition plays a major role in sensitization of the channel upon phospholipase C (PLC) activation by pro-inflammatory receptor stimulation (Chuang et al., 2001).

Subsequent studies from different laboratories however unanimously demonstrated that PtdIns(4,5)P_2_ or its precursor PtdIns(4)P, applied directly to excised inside out patches, potentiate TRPV1 activity (Stein et al., 2006; Lukacs et al., 2007; Klein et al., 2008; Lukacs et al., 2013a; Poblete et al., 2014). MgATP also potentiated channel activity in excised patches via stimulated synthesis of endogenous PtdIns(4,5)P_2_ (Lukacs et al., 2013a). When the purified channel was incorporated into planar lipid bilayers, the capsaicin- and heat-induced activity of TRPV1 also depended on the presence of PtdIns(4,5)P_2_ or PtdIns(4)P, demonstrating a direct effect on the channel (Lukacs et al., 2013a; Sun and Zakharian, 2015). A binding site responsible for the activating effect of PtdIns(4,5)P_2_ was proposed, based on the binding modes predicted by molecular docking of the lipid against the experimental cryoEM structure of TRPV1 (Poblete et al., 2014).

In intact cells, channel activity induced by capsaicin or low pH requires the presence of PtdIns(4,5)P_2_ and/or its immediate precursor PtdIns(4)P. Depletion of these lipids by specific inducible phosphoinositide phosphatases resulted in diminished channel activity (Klein et al., 2008; Yao and Qin, 2009; Hammond et al., 2012; Lukacs et al., 2013b). PtdIns(4,5)P_2_ and PtdIns(4)P are also depleted by Ca^2+^-induced activation of PLC when TRPV1 is activated by saturating capsaicin concentrations, and this effect plays an important role in desensitization of TRPV1 (Lishko et al., 2007; Lukacs et al., 2007; Yao and Qin, 2009; Lukacs et al., 2013b). These data provided overwhelming evidence for PtdIns(4,5)P_2_ being a positive cofactor/regulator of TRPV1, see more detailed discussion in (Rohacs, 2015).

When the purified TRPV1 channel was incorporated in lipid vesicles containing high concentrations of the negatively charged phosphatidylglycerol (~25%), its capsaicin sensitivity was reduced by PtdIns(4,5)P_2_, PtdIns(4)P_2_ or PtdIns, but not by PtdIns(3,4,5)P_3_ (Cao et al., 2013). Furthermore, the most recent and highest resolution cryoEM structure of TRPV1 revealed the presence of an endogenous phosphoinositide partially occupying the vanilloid binding site, and it was proposed that phosphoinositides act as competitive vanilloid antagonist (Gao et al., 2016). However this model is difficult to reconcile with the overwhelming evidence for PtdIns(4,5)P_2_ and PtdIns(4)P being positive regulators of TRPV1 in cellular membranes.

Here we attempt to resolve this controversy. Our data indicate that the endogenous lipid in the vanilloid binding site is phosphatidylinositol (PtdIns), the precursor of PtdIns(4)P and PtdIns(4,5)P_2_. We performed molecular dynamics (MD) simulations and found that PtdIns is more stable in the vanilloid binding site than PtdIns(4,5)P_2_. We showed that in excised patches PtdIns inhibited channel activity induced by low, but not by high capsaicin concentrations in the presence of PtdIns(4,5)P_2_, which is compatible with PtdIns acting as a competitive vanilloid antagonist. In the absence of PtdIns(4,5)P_2_, on the other hand, PtdIns partially stimulated TRPV1 activity especially in the presence of high capsaicin concentrations, an indication that PtdIns may also bind to the activating lipid binding site. We also found that mutating residues, predicted to interact with PtdIns, but not with capsaicin, resulted in higher sensitivity to capsaicin, as expected if PtdIns was a competitive antagonist. Our model resolves an important controversy and it is compatible with most data in the literature.

## RESULTS

### Two phosphoinositide binding sites in TRPV1

First, we compared the phosphoinositide binding site in TRPV1, which overlaps with the vanilloid pocket (Gao et al., 2016), with the one recently identified in the cryoEM structure of TRPV5 for PtdIns(4,5)P_2_ (Hughes et al., 2018). The PtdIns(4,5)P_2_ binding site in TRPV5 is strikingly similar to that proposed to be responsible for the activating effect of PtdIns(4,5)P_2_ in TRPV1 (Poblete et al., 2014). The TRPV5 structure was determined with diC_8_ PtdIns(4,5)P_2_ added to the purified protein, and the lipid induced a clear conformational change leading to channel opening (Hughes et al., 2018). The TRPV1 structure on the other hand was determined without adding a specific phosphoinositide; the lipid in the vanilloid binding site is likely derived from the soybean lipid extract added during the purification process, which contains a variety of phospholipids, including PtdIns (Gao et al., 2016). The lipid was modeled as PtdIns, but the authors argued that the site can accommodate a variety of further phosphorylated inositide lipids, including PtdIns(4,5)P_2_.

We thus superimposed the structures of TRPV1 (Gao et al., 2016) and TRPV5 (Hughes et al., 2018) to evaluate whether or not PtdIns and PtdIns(4,5)P_2_ binds to the same region of the channel. Comparison between the two structures clearly shows that the two binding sites are distinct: adjacent yet non-overlapping (**Figure 1; Figure S1**). The binding site of PtdIns overlaps with the vanilloid site, however it does not completely coincide with it; given the possible competition between PtdIns and capsaicin, hereafter we will refer to this site as “inhibiting” site. The binding site of PtdIns(4,5)P_2_ is completely consistent with the location previously proposed to mediate the positive regulatory role of PtdIns(4,5)P_2_ (Poblete et al., 2014); for this reason, hereafter we refer to this site as “activating site”. While the inhibiting site is located at the interface between the S1-S4 domain and the TRP-Domain, the activating site involves the S4-S5 linker, the C-terminus part of S6 and part of the TRP-Domain. (**Figure 1 and Figure S1**)

**Figure 1.**
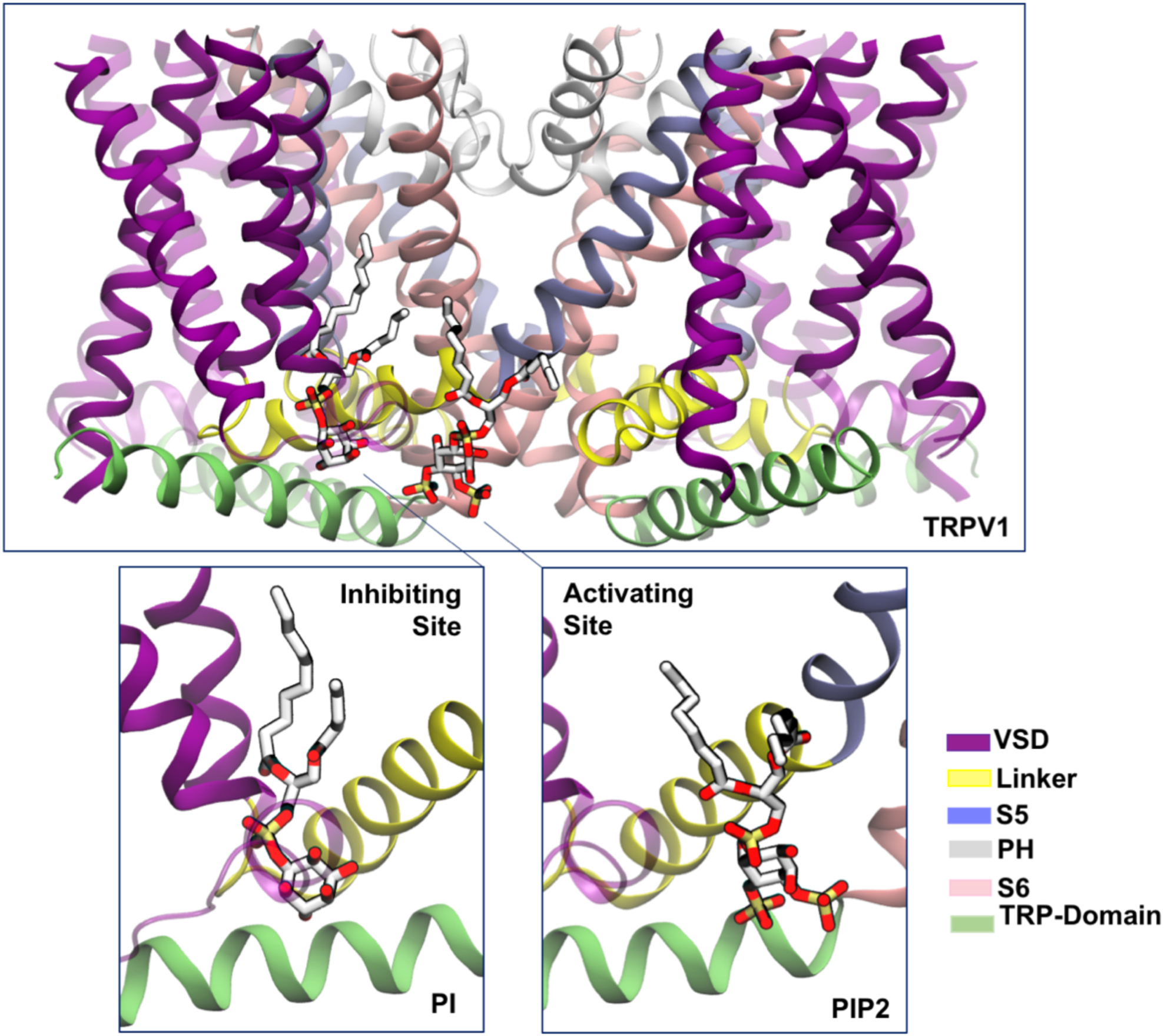
Phosphoinositides bind TRPV1 at two separate sites. Top panel. The transmembrane portion of the TRPV1 channel is shown (residues 430 to 713; side view), with phosphoinositides bound at the inhibiting (left) and activating (right) sites. The inhibiting site corresponds to the location of the phosphoinositide solved by Cryo-EM (Gao et al., 2016), and partially overlaps (but doesn’t coincide) with the vanilloid pocket; the activating site matches the experimentally observed location for the binding of 4,5-bisphosphate (PtdIns(4,5)P_2_ or PIP2) to TRPV5 (Hughes et al., 2018). The inhibiting and activating binding sites accommodate PtdIns and PtdIns(4,5)P_2_, respectively. The binding sites are shown for one subunit only using a different color scheme for each domain of the channel: the S1-S4-helix bundle is purple (residues 430 to 556), the S4-S5 linker is yellow (residues 557 to 575), S5 is ice-blue (residues 576 to 598), the pore helix (PH) is white (residues 599 to 655), S6 is pink (residues 656 to 689) and TRP-Domain is green (residues 690 to 713). In the insets, phosphoinositides atoms are shown as balls and sticks, color coded by element: C, O and P atoms are grey, red and yellow, respectively; hydrogen atoms not shown.

### The inhibiting site favors PtdIns, the activating site favors PtdIns(4,5)P_2_

As PtdIns(4,5)P_2_ and PtdIns(4)P are well-established positive regulators of TRPV1, we set out to test the hypothesis that the phosphoinositide occupying the vanilloid binding site is PtdIns. To this end we performed molecular dynamics (MD) simulations of TRPV1 with PtdIns (PI) or PtdIns (4,5)P_2_ (PIP2) in the putative inhibiting binding site. When we compared the root-mean-square deviation (RMSD) of atomic positions of each phosphoinositide along the two MD trajectories (0.42 μs each), we found that PtdIns is significantly more stable than PtdIns (4,5)P_2_ (**Figure 2 A** and **B**, respectively). To further corroborate the hypothesis that the inhibiting binding site favors PtdIns, we calculated the electronic density maps of PtdIns and PtdIns (4,5)P_2_ predicted by our simulations and compared them with the experimental density map (**Figure 2 E-H)**. PtdIns provides a better fit between simulation and experiment compared to PtdIns (4,5)P_2_.

**Figure 2.**
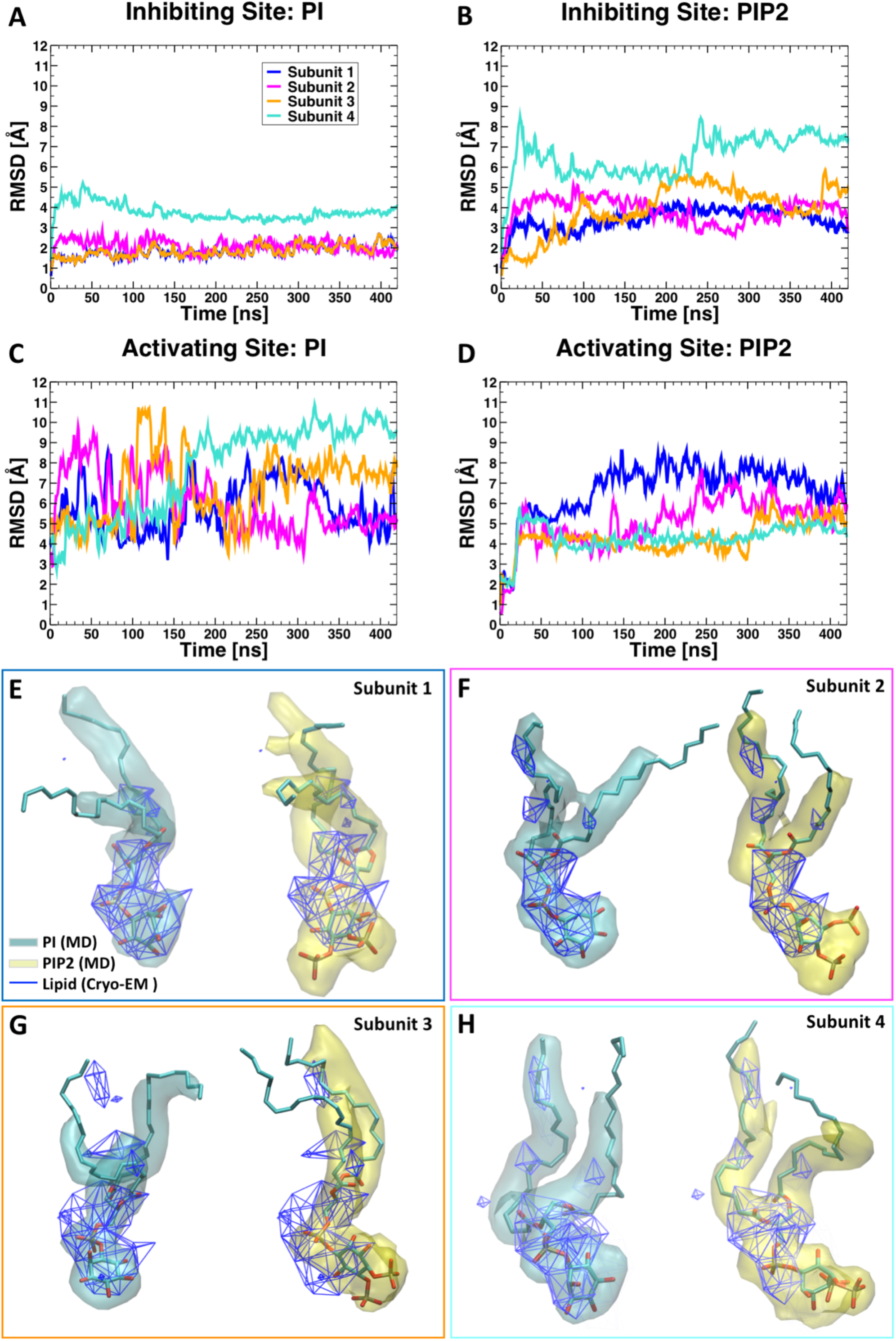
Structural stability of PtdIns and PtdIns(4,5)P_2_ at the inhibiting and the activating sites. (A)-(D). Root-mean-square deviation (RMSD) with respect to the initial configuration as a function of time from molecular dynamics simulations (a different color is used for each of the four symmetry-related subunits). Panels A and B show the RMSD of PtdIns and PtdIns(4,5)P_2_, respectively, when bound at the inhibiting site. Panels C and D show the RMSD of PtdIns and PtdIns(4,5)P_2_, respectively, when bound at the inhibiting site. (E)-(H). Superposition between the experimental and calculated lipid densities. Individual subunits are compared separately; each panel is color coded according to panel A legend. Calculated densities for PtdIns and PtdIns(4,5)P_2_ are colored in cyan and yellow, respectively, while the experimental electron density maps are shown as a blue-colored wireframe.

We also performed a set of similar simulations with PtdIns or PtdIns(4,5)P_2_ bound at the activating site (two simulations, 0.42 μs each). Analysis of RMSD values (**Figure 2 C-D)** clearly shows that while PtdIns(4,5)P_2_ is quite stable in this binding site, PtdIns shows large fluctuations. We interpret this behavior as possible indication of a greater affinity of this site for PtdIns(4,5)P_2_ compared to PtdIns.

Overall our computational findings suggest that two phosphoinositide binding sites exist in the TRPV1 channel; one, the inhibiting site, is mostly overlapping with the vanilloid pocket and binds primarily PtdIns. The other, the activating site, may favor binding of PtdIns(4,5)P_2_ over PtdIns.

### PtdIns partially inhibits TRPV1 in excised patches in the presence of PtdIns(4,5)P_2_

If indeed the phosphoinositide in the vanilloid binding site is PtdIns, then this lipid is expected to inhibit capsaicin-induced channel activity. To test this, we expressed TRPV1 in Xenopus oocytes, and performed excised inside out patch clamp experiments. We found that TRPV1 activity shows a decrease (run down) after patch excision in the presence of 0.5 μM capsaicin in the patch pipette (**Fig. 3**), which is consistent with our earlier findings (Lukacs et al., 2013a). This rundown is a characteristic of PtdIns(4,5)P_2_ dependent ion channels, caused by dephosphorylation of PtdIns(4,5)P_2_ by phosphatase enzymes in the patch membrane. When we applied the water soluble dioctanoyl (diC_8_) PtdIns(4,5)P_2_ (50 μM) channel activity was restored. Co-application of 50 μM diC_8_ PtdIns induced a partial, reversible inhibition of TRPV1 activity (**Fig. 3A**). Next we tested if the inhibitory effect of PtdIns can be decreased or eliminated by higher capsaicin concentrations, as expected if the lipid acts as a competitive vanilloid antagonist. **Figure 3C** shows that in the presence 4 μM capsaicin in the patch pipette, 50 μM PtdIns did not inhibit TRPV1 activity evoked by 50 diC_8_ PtdIns(4,5)P_2_.

**Figure 3.**
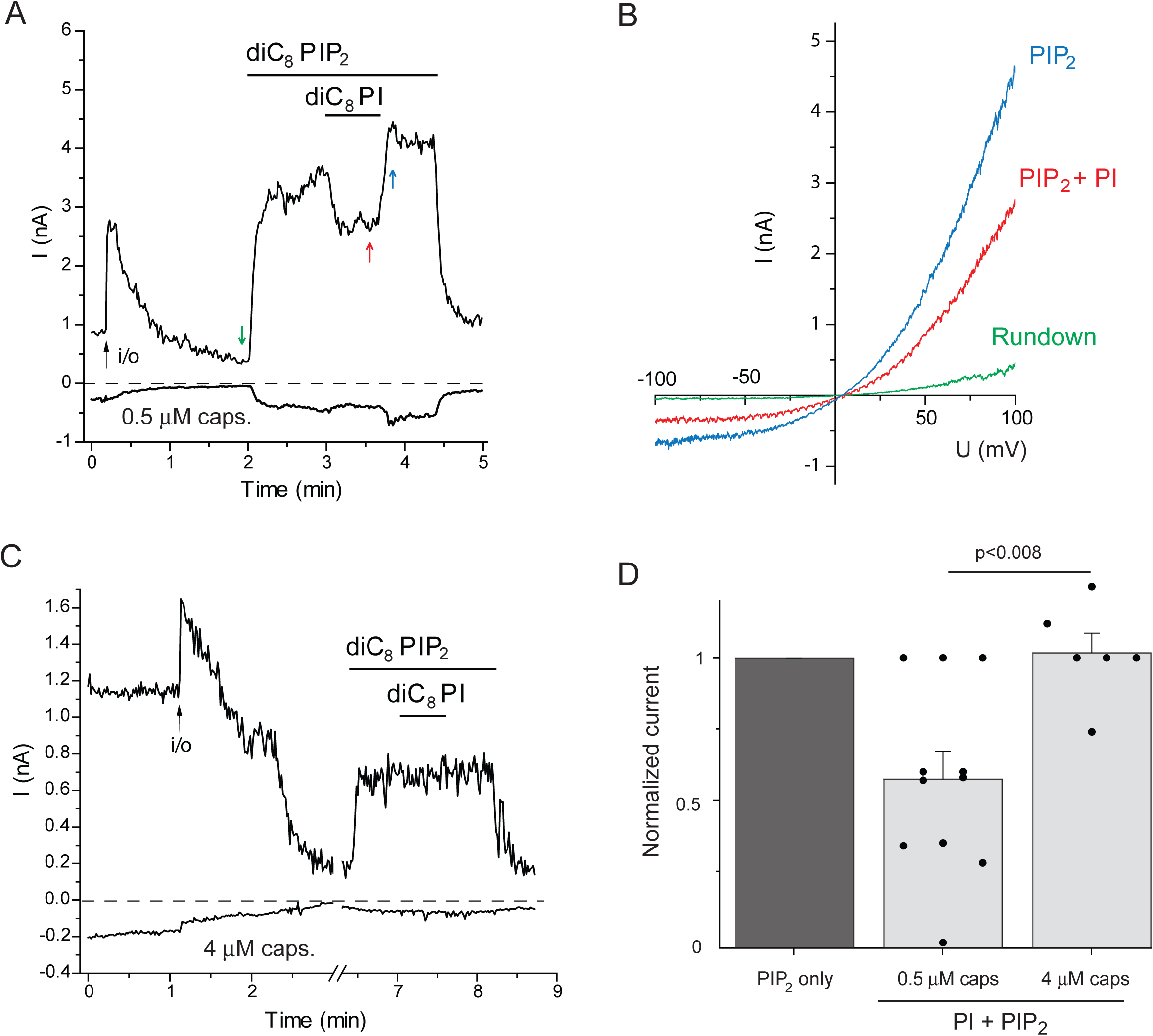
Phosphatidylinositol inhibits TRPV1 in excised patches at a low but not at high capsaicin concentration in the presence of PtdIns(4,5)P_2_. Excised inside out patch clamp experiments were performed as described in the Methods section in Xenopus oocytes injected with the TRPV1 cRNA. (A). Representative current trace at −100 and 100 mV, in the presence of 0.5 μM capsaicin in the patch pipette, the establishment of the inside out configuration is marked with an arrow, the applications of 50 μM diC_8_ PtdIns(4,5)P_2_ and 50 μM diC_8_ PtdIns are shown with the horizontal line. (B). Individual ramp current traces at the time points indicated by the colored arrows from panel A. (C). Representative current trace at −100 and 100 mV, in the presence of 4 μM capsaicin in the patch pipette, the establishment of the inside out configuration is marked with an arrow, the applications of 50 μM diC_8_ PtdIns(4,5)P_2_ and 50 μM diC_8_ PtdIns are shown by the horizontal lines. (D). Summary of the data, mean ± S.E.M. and scatter plots.

### PtdIns partially activates TRPV1 in the absence of PtdIns(4,5)P_2_

We propose here that PtdIns is a channel inhibitor at the binding site mostly overlapping with that for vanilloids. We found earlier that after channel rundown very high concentrations (500 μM) of long acyl chain (AASt) PtdIns partially reactivated TRPV1 at 0.5 μM capsaicin (Lukacs et al., 2013a). Here we revisited this and tested the effect of diC_8_ PtdIns at different capsaicin concentrations. We found that 100 μM diC_8_ PtdIns had no effect in the majority of patches in the presence of 0.5 μM capsaicin (**Figure 4A,E**), even though in some patches it induced a partial activation, which was smaller than that induced by 25 μM PtdIns(4,5)P_2_ (see **Figure 4B** for representative trace and **Figure 4E** for summary). In the presence of 4 μM capsaicin 100 μM diC_8_ PtdIns reliably reactivated TRPV1, but the effect was still smaller than that induced by 25 μM diC_8_ PtdIns(4,5)P_2_ (**Figure 4C-E**). These data indicate that capsaicin not only increases the apparent affinity for PtdIns(4,5)P_2_ in activating TRPV1 (Lukacs et al., 2007), but also decreases the selectivity of the activating phosphoinositide binding site allowing PtdIns to partially activate the channel.

**Figure 4:**
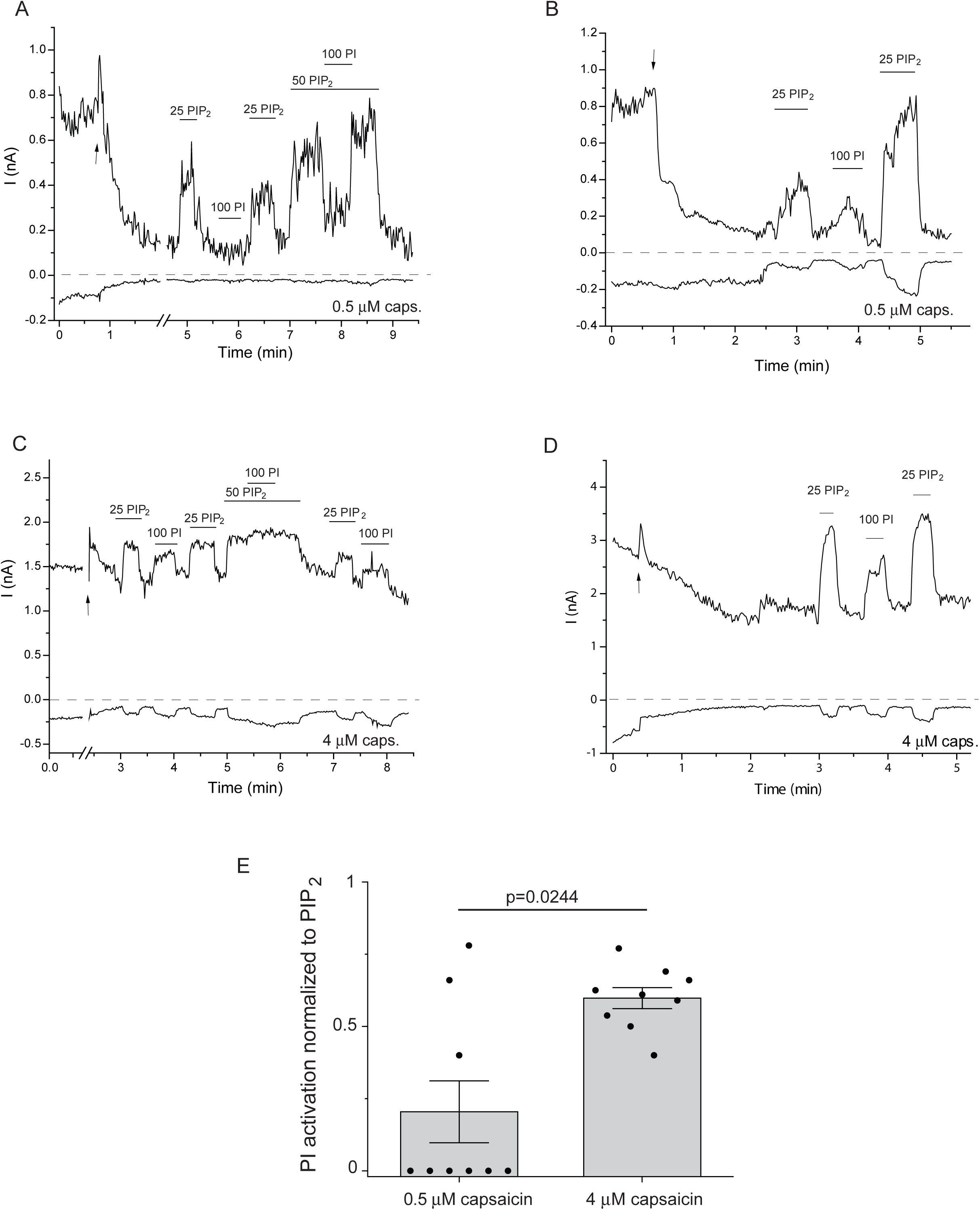
PtdIns partially activates TRPV1 in the presence of high capsaicin concentrations. Excised inside out patch experiments were performed as described in the Methods section in Xenopus oocytes injected with the TRPV1 cRNA. **(A)**, (**B**). Representative current traces at −100 and 100 mV, in the presence of 0.5 μM capsaicin in the patch pipette, the establishment of the inside out configuration is marked with an arrow, the applications of 25 and 50 μM diC_8_ PtdIns(4,5)P_2_ and 50 and 100 μM diC_8_ PtdIns are shown with the horizontal line. **(C)**, **(D)** Representative current traces at −100 and 100 mV, in the presence of 4 μM capsaicin in the patch pipette, the establishment of the inside out configuration is marked with an arrow, the applications of 25 and 50 μM diC_8_ PtdIns(4,5)P_2_ and 50 and 100 μM diC_8_ PtdIns are shown with the horizontal line. (**E)** Summary of the data: mean ± S.E.M. and scatter plots.

### Mutations predicted to affect PtdIns but not vanilloid binding increase sensitivity to capsaicin

Next we computationally identified residues, which are in contact with PtdIns, but not with capsaicin in the inhibiting binding site (5 angstroms cutoff): R409 and H410 in pre-S1, D509 and S510 in the S2-S3 loop, K571 and L574 in the S4-S5 segment, and I696, L699, Q700 and I703 in the TRP box (**Figure 5A**).

**Figure 5:**
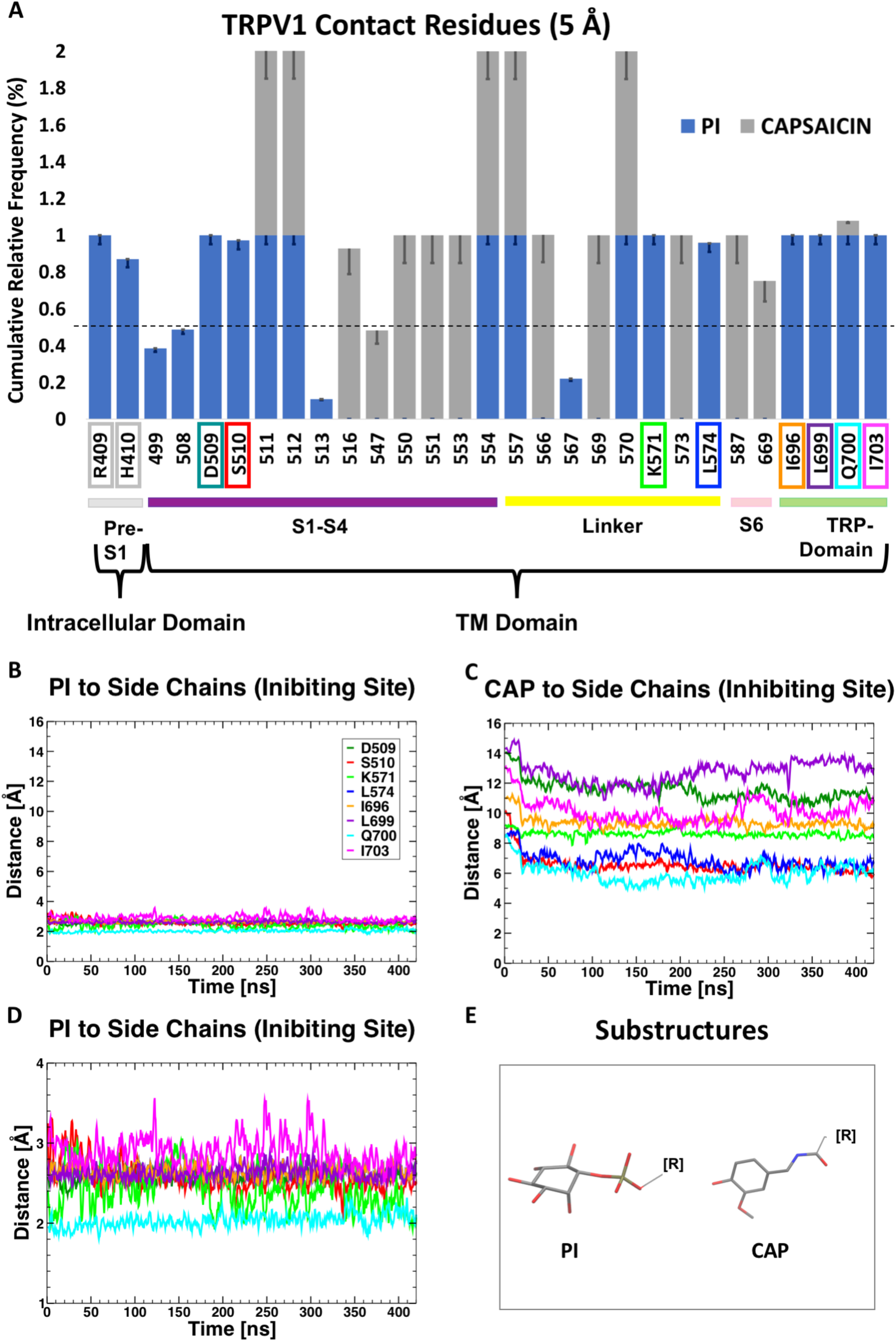
Residues in contact with PtdIns but not with capsaicin. **(A).** Analyses of residue contacts (from MD) identify what residues interact with PtdIns or with capsaicin. Not all the sidechains are in contact with both PtdIns and capsaicin, suggesting that the two binding sites (PtdIns and capsaicin) are in fact distinct. **(B)-(D).** Minimum distances between PtdIns (panels B and D) or capsaicin (panel E) and the sidechains shown in panel A. Distances are calculated between amino-acid side-chain atoms and lipid head groups (substructures in panel E), and averaged over all subunits. **(D).** Close-up of the time series shown in panel B.

In principle, mutation of a residue important for PtdIns binding but not for capsaicin binding, should reduce inhibition of the channel by endogenous PtdIns, and it would be predicted to increase sensitivity to activation by capsaicin (left shifted concentration response curve), because of weakening inhibition by an endogenous competitive inhibitor. We noticed that R409 and H410 are solvent exposed (**Figure 5 Supplement 1**) and, on average, more than 4 Angstroms farther away from PtdIns, therefore not in direct contact with it. Mutation of these residues is thus unlikely to produce large changes in PtdIns affinity. We thus focused our attention to D509, S510, K571, L574, I696, L699, Q700 and I703. All these sidechains are in close contact with PtdIns (with minimum distances smaller than 3 Angstroms, **Figure 5B,D**) yet are seemingly not interacting with capsaicin (minimum distances larger than 6 Angstroms, **Figure 5C**).

We thus considered these contacts as *bona fide* interactions and proceeded with a more quantitative (and computational intensive) characterization of these sidechains. In particular, we calculated the change in PtdIns affinity (ΔΔG) upon mutation of each sidechain into alanine. **Figure 6** shows that, with the exception of S510A, all the alanine mutants have a large destabilizing effect on PtdIns binding. Before proceeding with experimental testing of these mutants, we re-examined in detail the interactions between these sidechains and all the ligands involved (PtdIns(4,5)P_2_, PtdIns and capsaicin). We noticed that I696, K571 and L574 also form tight interactions with PtdIns(4,5)P_2_ in the activating site (**Figure 6 Supplement 1, panel A**) and therefore their mutation is likely to result in complex phenotypes. The same is true for Q700, which is involved in an indirect interaction with capsaicin through an intervening hydrogen-bonded water molecule (**Figure 6 Supplement 1, panel B**). Of the remaining list of amino acids, L699A had been reported to show a capsaicin sensitivity very similar to that of wild type channels (Valente et al., 2008).

**Figure 6:**
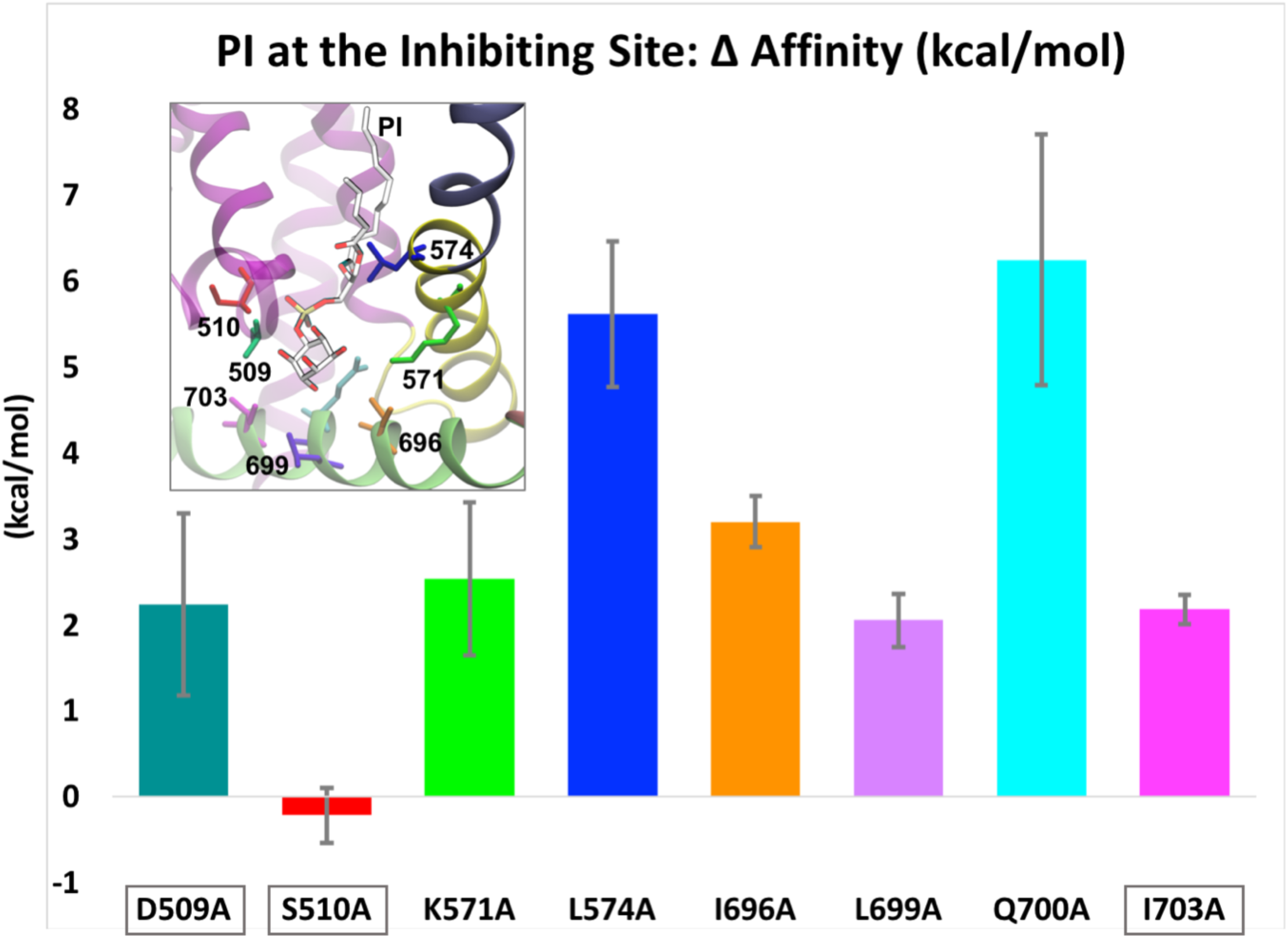
Change in affinity upon mutation of inhibiting binding site residues. The histogram plot shows the ΔΔG (kcal/mol), or change in PtdIns binding free energy, upon mutation into alanine of the binding site residues. Values are averaged over individual subunits and multiple configurations sampled from the MD trajectory. Error bars show the standard deviation. **Inset**. Binding mode of PtdIns at the inhibiting site. Residues selective for PtdIns binding, and subject to computational mutation analysis, are shown as sticks and colored according to the Legend in Fig.5.

For these reasons, we focused on residues D509 and I703 and mutated them individually to alanine. We also generated the S510 as a negative control. As shown in **Figure 7A-C** the D509A and I703A mutants showed left-shifted concentration dependence for capsaicin activation. The S510A mutant was somewhat less sensitive to capsaicin than the wild-type TRPV1 (data not shown). We further characterized the D509A and I703A mutants and we found that they also became more sensitive to activation by low pH (**Fig. 7 Figure Supplement 1**). This latter finding is not unexpected, as vanilloid antagonists such as capsazepine and iodinated resiniferatoxin inhibit TRPV1 responses not only to capsaicin, but also to low pH (Seabrook et al., 2002).

**Figure 7:**
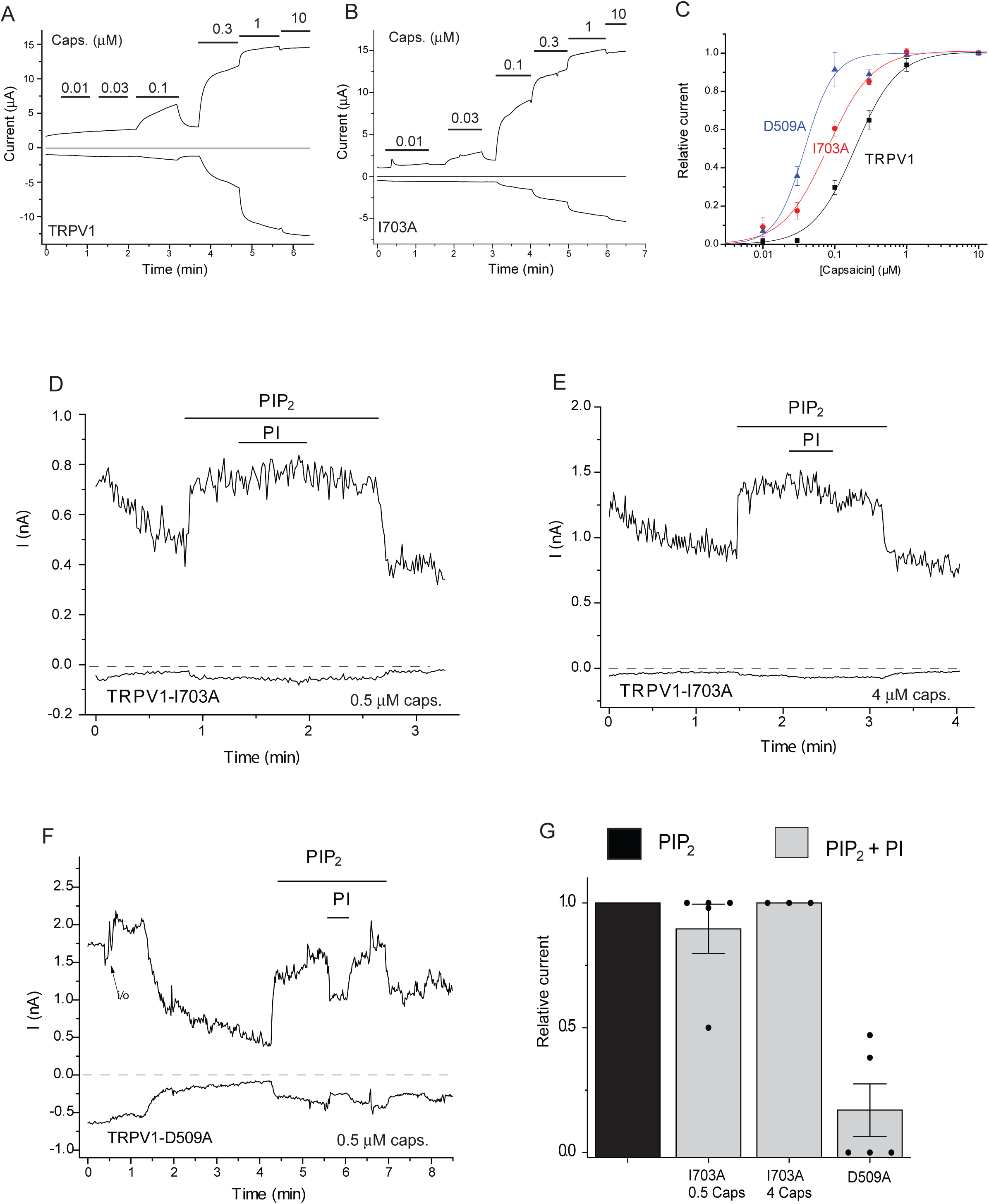
The I703A and D509A mutants are more sensitive to activation by capsaicin. **(A-C)** Two electrode voltage clamp experiments were performed as described in the Methods section in Xenopus oocytes injected with the TRPV1 and mutant cRNA. **(A-B)** Representative current traces at −100 and +100 mV the applications of different concentrations of capsaicin (μM) are indicated by the horizontal lines. **(C)** Hill plots for the concentration dependence of activation by capsaicin, error bars are S.E.M, n=7-12. **(D-F)**. Excised inside out patch experiments were performed as described in the Methods section in Xenopus oocytes injected with the TRPV1-I703A and D509A cRNA. **(D)** Representative current trace at −100 and 100 mV, in the presence of 0.5 μM capsaicin in the patch pipette the applications of 50 μM diC_8_ PtdIns(4,5)P_2_ and 50 μM diC_8_ PtdIns are shown with the horizontal lines. **(E)**. Representative current trace at −100 and 100 mV, in the presence of 4 μM capsaicin in the patch pipette, the applications of 50 μM diC_8_ PtdIns(4,5)P_2_ and 50 μM diC_8_ PtdIns are shown with the horizontal line. **(F)** similar measurement with the D509A mutant. **(G)** Summary of the data: mean ± S.E.M. and scatter plots.

Guided by our computational results, we then tested if these mutations also altered inhibition of TRPV1 by PtdIns in excised patches. **Figure 7D,E,G** shows that PtdIns did not inhibit the I703A mutant in excised patches neither in the presence of 0.5 μM capsaicin nor in the presence of 4 μM capsaicin. The D509A mutant on the other hand was inhibited by PtdIns (**Fig. 7F,G**), therefore the alteration of the capsaicin concentration response relationship was probably due to secondary effects unrelated to PtdIns. Our data with the I703A mutation on the other hand gives support to PtdIns inhibiting the channel via a binding site overlapping with that for vanilloid compounds such as capsaicin.

## DISCUSSION

The major goal of this work was to determine the nature of the endogenous phosphoinositide in the vanilloid binding site of TRPV1 (Gao et al., 2016), and to reconcile seemingly controversial data in the literature. Overall we conclude that the phosphoinositide occupying the vanilloid binding site is PtdIns, and we show that this lipid partially inhibits TRPV1 activity induced by low concentrations of capsaicin in the presence of PtdIns(4,5)P_2_. PtdIns(4,5)P_2_, the well-established positive cofactor, on the other hand activates the channel via a distinct, non-overlapping binding site, computationally predicted earlier by (Poblete et al., 2014). The latter binding site is similar to the one occupied by PtdIns(4,5)P_2_ in the structure of TRPV5 (a closely related channel in the TRP family) recently determined by cryoEM (Hughes et al., 2018). At saturating capsaicin concentrations PtdIns does not inhibit channel activity, as expected from a competitive inhibitor. PtdIns may also bind to the PtdIns(4,5)P_2_ binding site and partially activates the channel thus potentially exerting a dual effect.

In resting conditions, PtdIns(4,5)P_2_ constitutes ~1% of the plasma membrane phospholipids, and this lipid is a substrate for PLC enzymes (Balla, 2013). Its immediate precursor PtdIns(4)P is found at similar concentrations, and it can also be hydrolyzed by PLC, even though less efficiently than PtdIns(4,5)P_2_. Most research on ion channel regulation focused on PtdIns(4,5)P_2_ (Suh and Hille, 2008), but distinct roles for PtdIns(4)P have also been proposed (Hammond et al., 2012; Lukacs et al., 2013b).

PtdIns, the precursor of PtdIns(4,5)P_2_ and PtdIns(4)P, is thought to be at higher concentrations in the plasma membrane (up to 10%) but very little is known about its effect on ion channel function (Balla, 2013). Our molecular dynamics (MD) simulations indicate that PtdIns is more stable in the vanilloid binding site than PtdIns(4,5)P_2_, thus PtdIns most likely occupies this binding site. We found that applying PtdIns directly to excised patches partially inhibited channel activity induced by PtdIns(4,5)P_2_ in the presence of a low capsaicin concentration (0.5 μM). In the presence of a saturating concentration of capsaicin (4 μM) PtdIns exerted no inhibitory effect, as expected from a competitive antagonist.

Our computational data also show that PtdIns(4,5)P_2_ is more stable in the activating binding site than PtdIns. Consistent with this, PtdIns did not activate TRPV1 after current rundown in most patches in the absence of PtdIns(4,5)P_2_ at low (0.5 μM) capsaicin, even though it induced a small partial activation in some patches. In the presence of saturating capsaicin (4 μM), PtdIns reliably activated TRPV1, but the effect was still smaller than that induced by PtdIns(4,5)P_2_. Overall we propose that PtdIns has a dual effect on TRPV1 activity; it binds to the vanilloid binding site serving as a competitive inhibitor, but it may also bind to the activating lipid binding site, especially at high capsaicin concentrations. The TRPV1 structure with PtdIns was determined with the addition of soybean polar lipids which contains 18.4 % PtdIns (avantilipids.com/product/541602) (Gao et al., 2016). Both our experimental and computational data suggest that the binding of PtdIns to the activating site is likely to be much weaker than that of PtdIns(4,5)P_2_, explaining why this lipid did not show up in the putative activating binding site in the TRPV1 nanodiscs structure (Gao et al., 2016).

We identified residues in the vanilloid binding site that are predicted to interact with PtdIns but not with capsaicin. We found that the D509A and I703A mutants shifted the capsaicin dose response to the left, as expected if PtdIns is a competitive antagonist of capsaicin binding. The mutant channels also became more sensitive to low pH, which is compatible with PtdIns not only acting as a competitive vanilloid antagonist, but also as a negative allosteric regulator. This is in line with phosphoinositides also reducing heat sensitivity in lipid vesicles (Cao et al., 2013) as well as vanilloid antagonists, such as capsazepine and iodoresiniferatoxin, inhibiting TRPV1 responses to heat and low pH (Seabrook et al., 2002).

The I703A mutation also eliminated inhibition by PtdIns in excised patches, but the D509A mutant was inhibited by PtdIns to similar extent to wild type TRPV1, indicating that the latter may have altered PtdIns sensitivity by a different mechanism. Alternatively, PtdIns may also inhibit the mutant channel as a partial agonist acting on the activating PtdIns(4,5)P_2_ binging site. In accordance with this, we found earlier that even very high concentrations of long acyl chain natural PtdIns (500 μM) only induced a smaller effect than 20 μM long acyl chain natural PtdIns(4,5)P_2_ (Lukacs et al., 2013a).

While no negative effect of PtdIns(4,5)P_2_ was reported in excised patches, the purified TRPV1 incorporated into lipid vesicles containing ~25 % of the negatively charged Phosphatidylglycerol (PG) showed reduced sensitivity to capsaicin when various phosphoinositides were incorporated at 4%. PtdIns and PtdIns(4)P, induced the larger right shift in the capsaicin dose response than PtdIns(4,5)P_2_ (Cao et al., 2013). This order is the opposite of the effectiveness of phosphoinositides activating TRPV1 in excised patches where PtdIns(4,5)P_2_ and PtdIns(3,4,5)P_3_ had similar EC_50_ values (Klein et al., 2008), while PtdIns(4)P had a significantly lower apparent affinity, but similar maximal effect (Lukacs et al., 2007; Klein et al., 2008), whereas PtdIns had either no effect, or smaller than that induced by PtdIns(4,5)P_2_, current study and (Lukacs et al., 2013a). These data are compatible with our two binding site model, namely an inhibitory binding site favoring PtdIns, and an activating site preferring PtdIns(4,5)P_2_.

The inhibitory effect of PtdIns(4,5)P_2_ in lipid vesicles may also be attributed to incorporation of this lipid into the extracellular leaflet of the membrane, as it was shown that application of either PtdIns(4,5)P_2_ or PtdIns(4)P to the extracellular membrane surface in outside out patches inhibited TRPV1 currents (Senning et al., 2014). This effect however is unlikely to be mediated by the vanilloid binding site, which is located in the intracellular portion of the transmembrane regions of TRPV1. Also, PtdIns had no effect when applied to outside out patches (Senning et al., 2014), but inhibited TRPV1 when incorporated symmetrically into lipid vesicles (Cao et al., 2013).

When the purified TRPV1 was incorporated into planar lipid bilayers in a lipid mix without the negatively charged PG, channel activity induced by heat or capsaicin depended on the presence of PtdIns(4,5)P_2_ (Sun and Zakharian, 2015). PtdIns(4)P activated the channel less effectively than PtdIns(4,5)P_2_ at the same concentration, in agreement with excised patch data. When PG was incorporated in the lipid mix in the absence of PtdIns(4,5)P_2_ capsaicin opened the channels with an open probability close to 1, even though with a lower single channel conductance than that observed with PtdIns(4,5)P_2_ (Sun and Zakharian, 2015). In agreement with these data high concentrations of PG applied to excised patches, also partially reactivated TRPV1 after current rundown (Lukacs et al., 2013a). Overall, the most likely interpretation of these data is that TRPV1 requires a negatively charged lipid for activation by capsaicin. In a cellular environment this lipid is most likely PtdIns(4,5)P_2_ and to some extent PtdIns(4)P, but in artificial lipid mixtures, other lipids such as PG at high concentrations can also support channel activity.

The location of the PtdIns(4,5)P_2_ binding site in TRPV5 was recently determined experimentally (Hughes et al., 2018). TRPV5 is constitutively active in a cellular environment, but its activity depends on the presence of PtdIns(4,5)P_2_, and unlike in the case of TRPV1, no inhibitory effect has been proposed for this channel. Consistent with this, the TRPV5 structure in the presence of PtdIns(4,5)P_2_ showed a conformational change indicating clear channel opening, when compared to the structure of the channel without this lipid.

The location of the PtdIns(4,5)P_2_ binding site responsible for opening TRPV5 is very similar to that computationally predicted to be responsible for activation in TRPV1 (Poblete et al., 2014). Furthermore, we previously docked PtdIns(4,5)P_2_ to TRPV6 channels (Hughes et al., 2018), and the location of the lipid showed remarkable similarity to the one experimentally determined for the closely related TRPV5 (Hughes et al., 2018). These data provide convincing evidence that molecular docking of phosphoinositides against specific areas in the membrane cytoplasm interface can reliably predict the location of a lipid. By a similar approach, we used molecular docking to generate the starting configurations, for subsequent MD simulations, of PtdIns and PtdIns(4,5)P_2_ bound to TRPV1 in the vanilloid (inhibiting) or the activating sites, respectively. There configurations resulted to match almost perfectly with the experimental binding modes of lipids solved by cryoEM, whenever available (Gao et al., 2016). Our simulation results clearly show that PtdIns(4,5)P_2_ is more stable in the activating binding site than PtdIns, which is consistent with PtdIns(4,5)P_2_ being more efficient in opening the channel. While the final proof for the location of the activating phosphoinositide binding site in TRPV1 will likely come from future cryoEM studies with added PtdIns(4,5)P_2_, our two binding site model is compatible with most seemingly controversial data in the literature.

Overall, we identify PtdIns as a negative regulator of TRPV1 activity via the vanilloid binding site of the channel (the inhibiting site); this lipid may serve to set the basal sensitivity of the channel. We propose a two phosphoinositide binding site model, where the distinct non-overlapping activating binding site is mainly occupied by PtdIns(4,5)P_2_ in a cellular environment (**Figure 8**). Our data not only resolve a substantial controversy, but also identify a novel, specific role for PtdIns in regulating ion channel activity.

**Figure 8.**
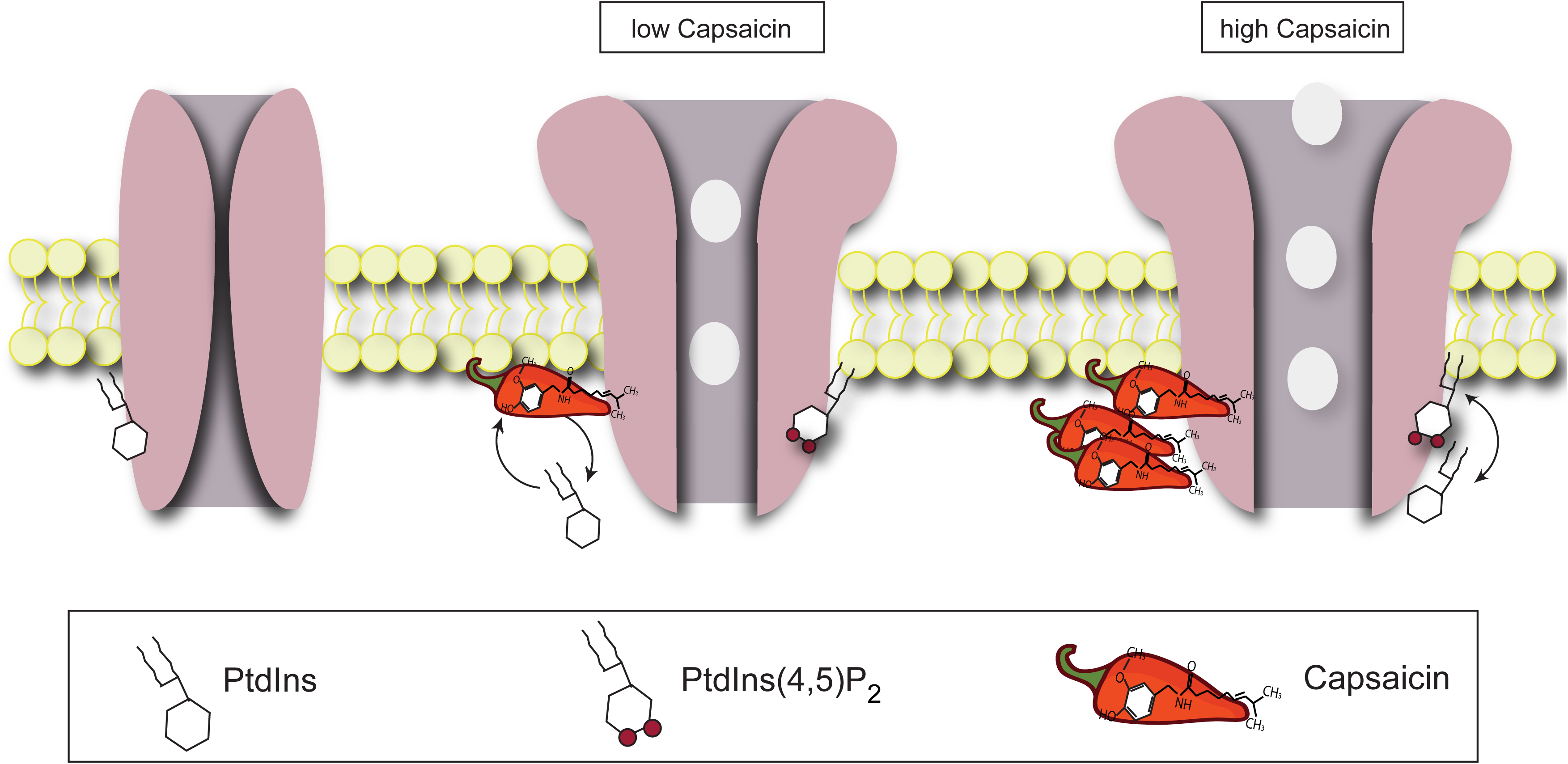
Cartoon explaining our two binding site model. **(left)** In the absence of capsaicin the vanilloid biding site is occupied by PtdIns. **(middle)** In the presence of low/moderate capsaicin concentrations capsaicin displaces PtdIns from the Vanilloid binding site and activates the channel, PtdIns(4,5)P_2_ in the activating binding site facilitates this activation. **(right)** In the presence of high capsaicin concentrations the vanilloid binding site is fully occupied by capsaicin, and both PtdIns and PtdIns(4,5)P_2_ can bind to the activating site; the channel is fully open.

## METHODS

### Xenopus Laevis oocyte preparation

Animal procedures were approved by the Institutional Animal Care and Use Committee at the New Jersey Medical School, and all animal procedures were performed in accordance with the approved guidelines. Oocytes were prepared from female *Xenopus laevis* frogs, as described earlier (Rohacs, 2013). Briefly: oocytes were digested using 0.2 mg/ml collagenase (Sigma) in a solution containing 82.5 mM NaCl, 2 mM KCl, 1 mM MgCl_2_, and 5 mM HEPES, pH 7.4 (OR2) overnight for ~16 h at 18 °C in a temperature controlled incubator. Defolliculated oocytes were selected and maintained in OR2 solution supplemented with 1.8 mM CaCl_2_ and 1% penicillin/streptomycin (Mediatech) at 18 °C.

### Excised inside-out patch clamp measurements in Xenopus oocytes

Excised inside-out patch clamp measurements were performed as described earlier (Lukacs et al., 2013a). Briefly, cRNA of TRPV1 transcribed from the rat TRPV1 clone in the pGEMSH vector using the mMessage mMachine kit (Thermo Fisher) was injected into *Xenopus laevis* oocytes. Measurements were performed 5-7 days after injection, using borosilicate glass pipettes (World Precision Instruments) of 0.4–0.7 MΩ resistance, filled with a solution containing 96 mM NaCl, 2 mM KCl, 1 mM MgCl_2_, and 5 mM HEPES, (pH=7.4) supplemented with 0.5 μM or 4 μM capsaicin (as indicated in figure legends). After establishing GΩ-resistance seals on devitellinized surfaces of oocytes, inside-out configuration was established, and currents were measured using a ramp protocol from −100 to +100 mV applied every second. The main perfusion solution contained 96 mM KCl, 5 mM EGTA, and 10 mM HEPES, pH adjusted to 7.4. Currents were recorded with an Axopatch 200B unit and analyzed with the pClamp 9.2 software (Molecular Devices). Measurements were performed at 18– 20°C. Various stimulating solutions were applied to the internal side of the inside-out membrane patch using a custom-made gravity-driven perfusion system. DiC_8_ phosphoinositides were purchased from Cayman Chemical (Ann Arbor, MI). Capsaicin was purchased from Sigma (St. Louis, MO).

### Two-electrode voltage clamp (TEVC) in Xenopus oocytes

Measurements were performed as described earlier (Lukacs et al., 2007). Briefly, oocytes were placed in a solution containing 97 mM NaCl, 2 mM KCl, 1 mM MgCl_2_, and 5 mM HEPES, pH 7.4, currents were recorded with thin-wall inner filament-containing glass pipettes (World Precision Instruments) filled with 3 M KCl in 1% agarose. Currents were measured with an Axopatch 200B amplifier and analyzed with the pClamp 9.2 software (Molecular Devices) using the same ramp protocol as described earlier for excised patch measurements. Various stimulating solutions were applied using a gravity-driven perfusion system.

### Data analysis

Data are represented as mean ± S.E. plus scatter plots; statistical significance is calculated either with *t*-test, analysis of variance, or Mann-Whitney test as appropriate; *, p<0.05; **, p<0.01; ***, p<0.001

### Computational Methods

We used the structure of TRPV1 in lipid nanodiscs solved in the presence of a phosphoinositide at the vanilloid binding site (apo TRPV1; PDB-ID: 5irz) (Gao et al., 2016) as the starting configuration of all our computational studies. Since the experimental structural model contains only a truncated version of the phospholipid tails and the stereochemistry of the sugar head group does not match any of the naturally occurring isomers of myo-inositol phosphate, we reconstructed, refined and equilibrated the structure of the experimentally determined lipid. We used the charge and tautomeric states defined in (Jo et al., 2008). We thus docked the equilibrated structure of the refined lipid against the vanilloid binding site of TRPV1 (while preserving the experimental binding mode), before minimizing the obtained structural complex.

#### Molecular Docking

We used the docking tool Glide (Friesner et al., 2004) (Schrödinger, LLC, New York, NY, 2018) to refine the binding mode generated manually for PI against the inhibiting site of TRPV1. To this purpose, we started from the equilibrated structure of the channel (as described in the previous paragraph), which was properly prepared in the Maestro tool using the Protein Preparation Wizard (Sastry et al., 2013). The docking grid was centered around the manually docked configuration of PI. The structure of PI was prepared using LigPrep. Docking poses were prioritized by score (kcal/mol) and postprocessing was performed by cluster analysis, using the tool g_cluster with the Jarvis Patrick methodology available in GROMACS (Pronk et al., 2013; Abraham et al., 2015) (http://www.gromacs.org). The top 4 binding modes (one each TRPV1 subunit) were selected to constitute our final structural complex of TRPV1 with PI bound at the inhibiting site. In addition, we modified the structures of PI lipids in this final complex to match the geometry of PIP2, and by applying topology and atom types in agreement with the force field available through the CHARMM-GUI interface (SAPI25) (Jo et al., 2008). An analogous procedure was followed for the docking of PIP2 to the activating site.

#### Molecular Dynamics (MD) Simulations

We embedded the structures of TRPV1 in complex with phosphoinositides in a hydrated 1-palmitoyl-2-oleoylphosphatidylcholine (POPC) bilayer, using the membrane plugin of VMD (Humphrey et al., 1996). The system was then surrounded by 150 mM NaCl solution to reach an overall size of ~ 160×160×150 Å^3^, with a total number of 330,000 atoms. We used all-atom molecular dynamics (MD) simulation to equilibrate the systems (five in total: TRPV1 bound to PI or PIP2 at the inhibiting or the activating sites plus a TRPV1 in complex with capsaicin) through a multistep protocol. First, we performed energy minimization of the systems (1,000 steps). Second, we applied position restraints on protein (backbone and side chain atoms) and lipids (head groups) that were gradually released during the first 50-ns simulation time (harmonic potentials with initial force constant K1 = 20 kcal/mol/Å^2^ were applied to all restrained atoms). Last, upon releasing the restraints, we performed additional ~425-ns production runs. During all simulations, we used the velocity Verlet integration method to solve the equations of motion, with a time step of 2 fs using the Particle mesh Ewald (PME) method for calculating the electrostatic potential. We applied the Langevin temperature and Langevin piston coupling schemes. We set the temperature to 300 K and the pressure to 1 atm. We generated four independent MD trajectories, with PI or PIP2 bound at the inhibiting or the activating sites. In all cases, we used the CHARMM36 force field (Mackerell et al., 2004) to describe the protein and the POPC lipids, and the parameters derived from ref (Kasimova et al., 2018). and ref. (Jo et al., 2008) for capsaicin and phosphoinositide lipids, respectively. The TIP3P model was used to describe water molecules (Jorgensen et al., 1983). We used the VMD (Humphrey et al., 1996) (version 1.9) and NAMD (Phillips et al., 2005) (version 2.12) programs for system preparation, equilibration, MD simulation and trajectory analysis.

#### Isolation of Lipid Densities (Cryo-EM Map)

We used the software Chimera (version 1.13.1; https://www.cgl.ucsf.edu/chimera/) to isolate densities of phosphoinositide lipids occupying the inhibiting site in the TRPV1 structure solved by Cryo-EM (PDB-ID: 5irz). Specifically, the Cryo-EM map was uploaded on the experimental structure file and the “Volume Viewer” and the “Volume Eraser” tools were used to extract the density blobs of bound lipids, which were saved as BRIX files. The experimental densities of lipids were then compared by superimposition with atomic densities derived from MD simulations.

#### Atomic Density Maps Calculations (MD Simulations)

We calculated the atomic density maps for the phosphoinositide lipids (PI or PIP2) occupying the inhibiting or the activating sites using the VolMap Plugin available in VMD(9) (version 1.9). For all calculations, we used a resolution of 0.5 Å, with an atom size of 1.0 Å and weights corresponding to the atomic mass. Maps were computed for all frames of each trajectory and subunit; frames were then combined by the averaging method. The computed atomic densities of lipids were then compared by superimposition with the experimental densities obtained by Cryo-EM.

#### Analyses of Contact Residues

We first used custom script (Tcl) to select protein side-chain atoms within a distance threshold (5 Å) of headgroups in either the phosphoinositide lipid PI or capsaicin (CAP). Headgroups were defined as all heavy atoms in the substructures depicted in **Fig. 5E** (bold licorice). Residue contact lists, with relative frequency of occurrence within the distance threshold, were then generated individually per subunit, and finally combined over each simulation trajectory (TRPV1 with PI or CAP at the inhibiting site). Last, cumulative relative frequencies were plotted as histograms, color-coded by ligand type (PI is blue and CAP is gray), using Microsoft Excel. By comparing the two distributions, we identified 8 protein side-chains that were selectively in contact with PI (but not with CAP) in at least 50% of the analyzed simulation frames. These residue, specifically D509, S510, K571, L574, I696, L699, Q700 and I703, are located in the inhibiting site of TRPV1.

#### Min-Distance Calculations

We calculated the minimum distance between the headgroups (**Fig. 5E**, bold licorice) of the phosphoinositide lipid PI or capsaicin (CAP) occupying the inhibiting site and the contact residues within 5 Å of PI. Distances were calculated using the “g_mindist” function available with the program GROMACS (http://www.gromacs.org). Distance plots were generated in XMGRACE (Vaught, 1996). Substructures were sketched using Sketch Tool available in Maestro (Schrödinger, LLC, New York, NY, 2018).

#### ΔΔG Calculations

We calculated the change in binding affinity of TRPV1 for PI upon mutation of each PI-contacting residue into alanine. To do so, we used the “Residue-Scanning and Mutation” protocol available in BioLuminate (Beard et al., 2013) (Schrödinger, LLC, New York, NY, 2018). All residues (D509, S510, K571, L574, I696, L699, I703) were mutated to alanine side-chains; upon mutation, protein side chains were minimized to optimize interactions with the bound lipid. Residue mutations and *ΔΔG* calculations were performed on configurations sampled from the MD trajectory at ~150-ns intervals. As a control, we repeated all calculations with capsaicin bound at the vanilloid site (one frame). Systems were prepared for the calculations using the Protein Preparation Wizard (Sastry et al., 2013) (Schrödinger, LLC, New York, NY, 2018). For each system (4 frames in total; 3 TRPV1 bound to PI and 1 TRPV1 bound to capsaicin), we performed independent calculations for each subunit and mutant (4 subunits x 8 mutants). We finally carried out a total of 32 calculations. Results were averaged over all subunits (**Fig. 6**). Affinity changes were plotted using Microsoft Excel. Error bars in the histogram plots indicate standard deviation.

## ACKNOWLEDGMENTS

This work was supported, by National Institutes of Health Grants NS055159 (to T.R.) and GM093290 (to T. R. and E.C.) and National Science Foundation Grant ACI-1614804 (E.C.). This research includes calculations carried out on Temple University’s HPC resources and thus was supported in part by the National Science Foundation through major research instrumentation grant number 1625061 and by the US Army Research Laboratory under contract number W911NF-16-2-0189 as well as National Institutes of Health Grant S10OD020095

---

**Figure 1 Figure Supplement 1.**
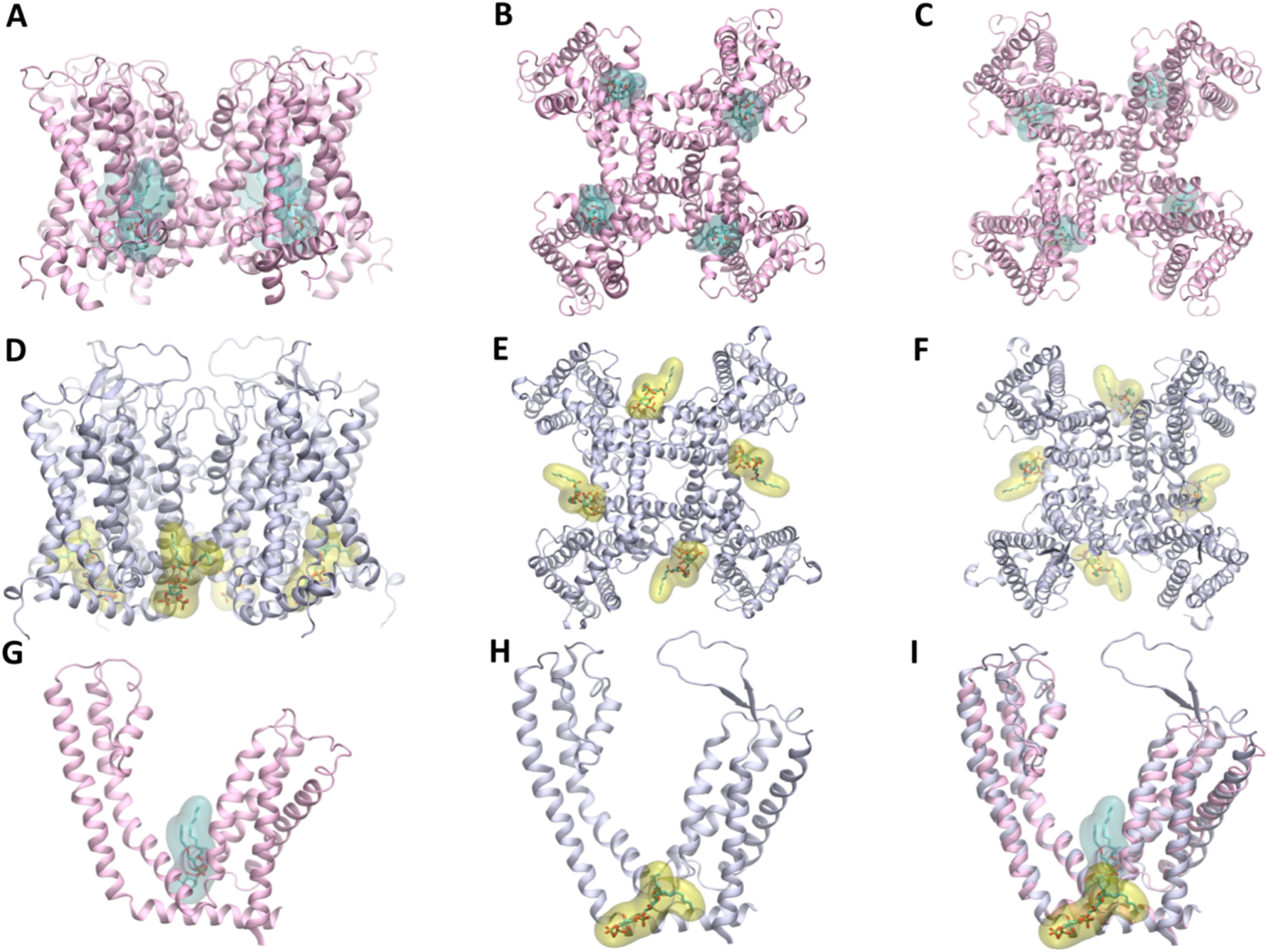
Binding of PtdIns and PtdIns(4,5)P_2_ at the inhibiting and the activating sites. (A)-(C). PtdIns bound at the inhibiting site of TRPV1 (Gao et al., 2016); side, top and bottom views, respectively. (D)-(F). PtdIns(4,5)P_2_ bound at the activating site of TRPV5 (Hughes et al., 2018); side, top and bottom view, respectively. (G). PtdIns bound at the inhibiting site of TRPV1; one subunit only is shown. (H). PtdIns(4,5)P_2_ bound at the activating site of TRPV5; one subunit only is shown. (I). Superimposition between PtdIns and PtdIns(4,5)P_2_ bound at the activating or inhibiting sites of TRPV1 or TRPV5, respectively. TRPV1 and TRPV5 are colored in pink and ice-blue, respectively. Phosphoinositides atoms are shown as balls and sticks, color coded by element: C, O and P atoms are grey, red and yellow, respectively; hydrogen atoms not shown. The molecular surfaces of PtdIns and PtdIns(4,5)P_2_ are also shown in cyan and yellow for PtdIns and PtdIns(4,5)P_2_, respectively.

**Figure 5 Figure Supplement 1:**
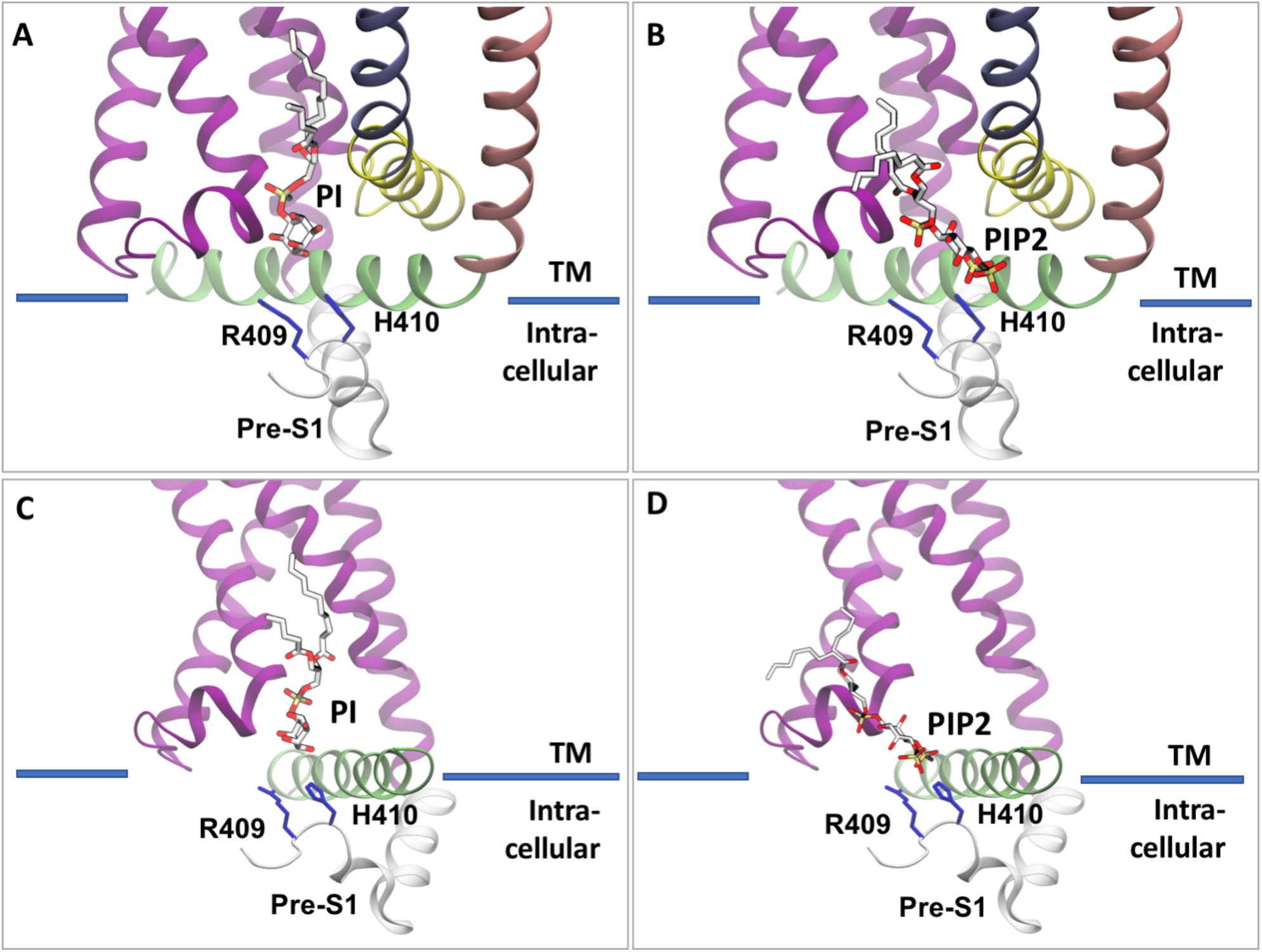
Two charged amino acids in contact with PtdIns are located on the pre-S1 segment. Among the residues selectively in contact with PtdIns over capsaicin, R409 and H410, in addition to being the only two charged amino acids, and are located on the pre-S1 segment, too far away to engage in stabilizing interactions with PtdIns. Localization of R409 and H410 with respect to PtdIns and PtdIns(4,5)P_2_ bound at the inhibiting **(A)-(C)** and the activating sites **(B)-(D)**, respectively.

**Figure 6 Figure Supplement 1.**
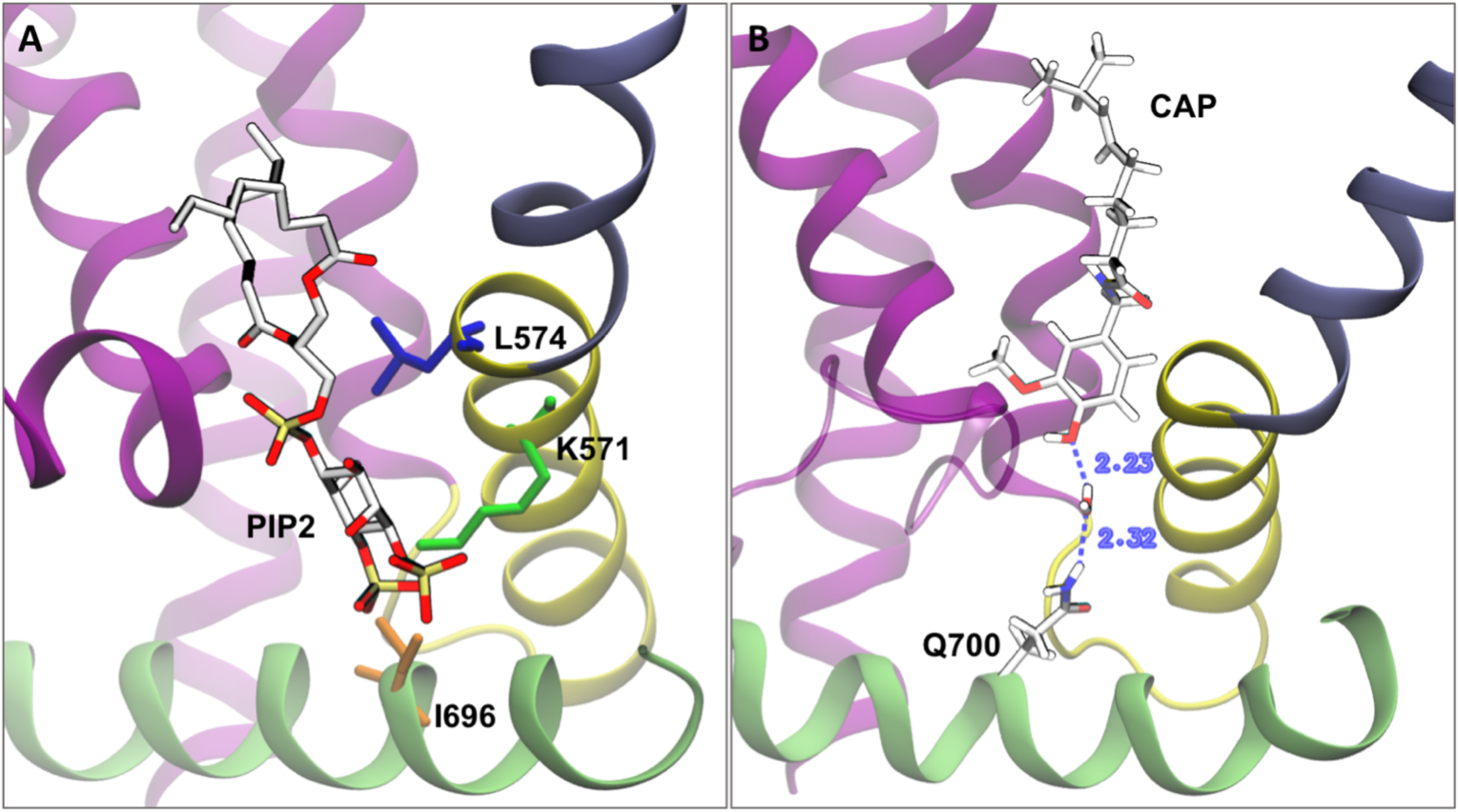
Side chains interacting with multiple ligands (PtdIns(4,5)P_2_, PtdIns and capsaicin). **(A)** Ball and sticks representation of the sidechains of L575, K571 and I696 and of PtdIns(4,5)P_2_ (note the close proximity). **(B)** Sidechain of residue Q700 observed in our molecular dynamics simulations to interact with capsaicin through an intervening hydrogen-bonded water molecule.

**Figure 7 Figure Supplement 1:**
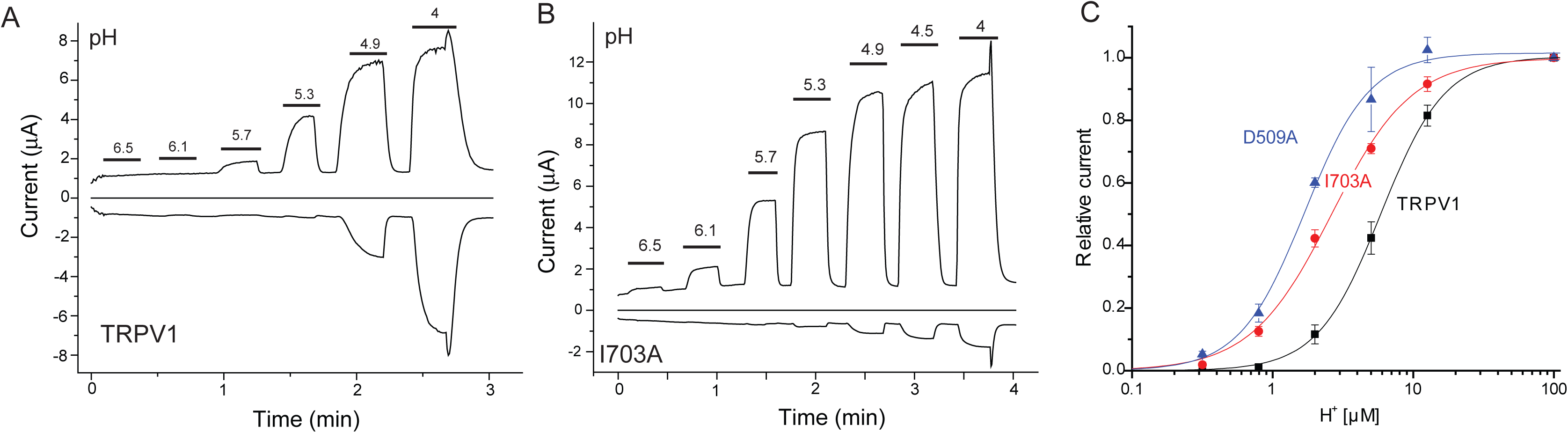
The I703A and D509A mutants are more sensitive to activation by low pH. Two electrode voltage clamp experiments were performed as described in the Methods section in Xenopus oocytes injected with the TRPV1 and mutant cRNA. **(A,B)** Representative current traces at −100 and +100 mV the applications of solutions with different pH are indicated by the horizontal lines. **(C)** Hill plots for the sensitivity of activation by low pH at 100 mV, expressed as [H^+^] in μM, error bars are S.E.M, n=5-7.

## REFERENCES

Abraham, M.J., Murtola, T., Schulz, R., Pall, S., Smith, J.C., Hess, B., and Lindahl, E. (2015). GROMACS: High performance molecular simulations through multi-level parallelism from laptops to supercomputers. SoftwareX 1-2, 19–25.

Balla, T. (2013). Phosphoinositides: tiny lipids with giant impact on cell regulation. Physiol Rev 93, 1019–1137.

Beard, H., Cholleti, A., Pearlman, D., Sherman, W., and Loving, K.A. (2013). Applying physics-based scoring to calculate free energies of binding for single amino acid mutations in protein-protein complexes. PLoS One 8, e82849.

Cao, E., Cordero-Morales, J.F., Liu, B., Qin, F., and Julius, D. (2013). TRPV1 channels are intrinsically heat sensitive and negatively regulated by phosphoinositide lipids. Neuron 77, 667–679.

Caterina, M.J., Leffler, A., Malmberg, A.B., Martin, W.J., Trafton, J., Petersen-Zeitz, K.R., Koltzenburg, M., Basbaum, A.I., and Julius, D. (2000). Impaired nociception and pain sensation in mice lacking the capsaicin receptor. Science 288, 306–313.

Caterina, M.J., Rosen, T.A., Tominaga, M., Brake, A.J., and Julius, D. (1999). A capsaicin-receptor homologue with a high threshold for noxious heat. Nature 398, 436–441.

Chuang, H.H., Prescott, E.D., Kong, H., Shields, S., Jordt, S.E., Basbaum, A.I., Chao, M.V., and Julius, D. (2001). Bradykinin and nerve growth factor release the capsaicin receptor from PtdIns(4,5)P2-mediated inhibition. Nature 411, 957962.

Friesner, R.A., Banks, J.L., Murphy, R.B., Halgren, T.A., Klicic, J.J., Mainz, D.T., Repasky, M.P., Knoll, E.H., Shelley, M., Perry, J.K., Shaw, D.E., Francis, P., and Shenkin, P.S. (2004). Glide: a new approach for rapid, accurate docking and scoring. 1. Method and assessment of docking accuracy. J Med Chem 47, 17391749.

Gao, Y., Cao, E., Julius, D., and Cheng, Y. (2016). TRPV1 structures in nanodiscs reveal mechanisms of ligand and lipid action. Nature 534, 347–351.

Hammond, G.R., Fischer, M.J., Anderson, K.E., Holdich, J., Koteci, A., Balla, T., and Irvine, R.F. (2012). PI4P and PI(4,5)P2 Are Essential But Independent Lipid Determinants of Membrane Identity. Science 337, 727–730.

Hughes, T.E.T., Pumroy, R.A., Yazici, A.T., Kasimova, M.A., Fluck, E.C., Huynh, K.W., Samanta, A., Molugu, S.K., Zhou, Z.H., Carnevale, V., Rohacs, T., and Moiseenkova-Bell, V.Y. (2018). Structural insights on TRPV5 gating by endogenous modulators. Nat Commun 9, 4198.

Humphrey, W., Dalke, A., and Schulten, K. (1996). VMD: visual molecular dynamics. J Mol Graph 14, 33–38, 27-38.

Jo, S., Kim, T., Iyer, V.G., and Im, W. (2008). CHARMM-GUI: a web-based graphical user interface for CHARMM. J Comput Chem 29, 1859–1865.

Jorgensen, W.L., Chandrasekhar, J., Madura, J.D., Impey, R.W., and Klein, M.L. (1983). Comparison of simple potential functions for simulating liquid water. J Chem Phys 79, 926.

Kasimova, M., Yazici, A., Yudin, Y., Granata, D., Klein, M., Rohacs, T., and Carnevale, V. (2018). Ion Channel Sensing: Are Fluctuations the Crux of the Matter? The Journal of Physical Chemistry Letters 9, 1260–1264.

Klein, R.M., Ufret-Vincenty, C.A., Hua, L., and Gordon, S.E. (2008). Determinants of molecular specificity in phosphoinositide regulation. Phosphatidylinositol (4,5)-bisphosphate (PI(4,5)P2) is the endogenous lipid regulating TRPV1. J Biol Chem 283, 26208–26216.

Lishko, P.V., Procko, E., Jin, X., Phelps, C.B., and Gaudet, R. (2007). The ankyrin repeats of TRPV1 bind multiple ligands and modulate channel sensitivity. Neuron 54, 905–918.

Lukacs, V., Rives, J.M., Sun, X., Zakharian, E., and Rohacs, T. (2013a). Promiscuous activation of transient receptor potential vanilloid 1 channels by negatively charged intracellular lipids, the key role of endogenous phosphoinositides in maintaining channel activity. J Biol Chem 288, 3500335013.

Lukacs, V., Thyagarajan, B., Varnai, P., Balla, A., Balla, T., and Rohacs, T. (2007). Dual regulation of TRPV1 by phosphoinositides. J Neurosci 27, 70707080.

Lukacs, V., Yudin, Y., Hammond, G.R., Sharma, E., Fukami, K., and Rohacs, T. (2013b). Distinctive changes in plasma membrane phosphoinositides underlie differential regulation of TRPV1 in nociceptive neurons. Journal of Neuroscience 33, 11451–11463.

Mackerell, A.D., Jr., Feig, M., and Brooks, C.L., 3rd (2004). Extending the treatment of backbone energetics in protein force fields: limitations of gas-phase quantum mechanics in reproducing protein conformational distributions in molecular dynamics simulations. J Comput Chem 25, 1400–1415.

Phillips, J.C., Braun, R., Wang, W., Gumbart, J., Tajkhorshid, E., Villa, E., Chipot, C., Skeel, R.D., Kale, L., and Schulten, K. (2005). Scalable molecular dynamics with NAMD. J Comput Chem 26, 1781–1802.

Poblete, H., Oyarzun, I., Olivero, P., Comer, J., Zuniga, M., Sepulveda, R.V., Baez-Nieto, D., Gonzalez Leon, C., Gonzalez-Nilo, F., and Latorre, R. (2014). Molecular Determinants of Phosphatidylinositol 4,5Bisphosphate (PI(4,5)P2) Binding to Transient Receptor Potential V1 (TRPV1) Channels. J Biol Chem 290, 2086–2098.

Pronk, S., Pall, S., Schulz, R., Larsson, P., Bjelkmar, P., Apostolov, R., Shirts, M.R., Smith, J.C., Kasson, P.M., van der Spoel, D., Hess, B., and Lindahl, E. (2013). GROMACS 4.5: a high-throughput and highly parallel open source molecular simulation toolkit. Bioinformatics 29, 845–854.

Rohacs, T. (2013). Recording macroscopic currents in large patches from Xenopus oocytes. Methods Mol Biol 998, 119–131.

Rohacs, T. (2014). Phosphoinositide regulation of TRP channels. Handb Exp Pharmacol 233, 1143–1176.

Rohacs, T. (2015). Phosphoinositide regulation of TRPV1 revisited. Pflugers Arch 467, 1851–1869.

Sastry, G.M., Adzhigirey, M., Day, T., Annabhimoju, R., and Sherman, W. (2013). Protein and ligand preparation: parameters, protocols, and influence on virtual screening enrichments. J Comput Aided Mol Des 27, 221–234.

Seabrook, G.R., Sutton, K.G., Jarolimek, W., Hollingworth, G.J., Teague, S., Webb, J., Clark, N., Boyce, S., Kerby, J., Ali, Z., Chou, M., Middleton, R., Kaczorowski, G., and Jones, A.B. (2002). Functional properties of the high-affinity TRPV1 (VR1) vanilloid receptor antagonist (4-hydroxy-5-iodo-3-methoxyphenylacetate ester) iodo-resiniferatoxin. J Pharmacol Exp Ther 303, 1052–1060.

Senning, E.N., Collins, M.D., Stratiievska, A., Ufret-Vincenty, C.A., and Gordon, S.E. (2014). Regulation of TRPV1 by Phosphoinositide (4,5)-bisphosphate: Role of Membrane Asymmetry. J Biol Chem 289, 10999–11006.

Stein, A.T., Ufret-Vincenty, C.A., Hua, L., Santana, L.F., and Gordon, S.E. (2006). Phosphoinositide 3-kinase binds to TRPV1 and mediates NGF-stimulated TRPV1 trafficking to the plasma membrane. J Gen Physiol 128, 509–522.

Suh, B.C., and Hille, B. (2008). PIP2 is a necessary cofactor for ion channel function: how and why? Annu Rev Biophys 37, 175–195.

Sun, X., and Zakharian, E. (2015). Regulation of the temperature-dependent activation of Transient Receptor Potential Vanilloid 1 (TRPV1) by phospholipids in planar lipid bilayers. Journal of Biological Chemistry 290, 4741–4747.

Valente, P., Garcia-Sanz, N., Gomis, A., Fernandez-Carvajal, A., Fernandez-Ballester, G., Viana, F., Belmonte, C., and Ferrer-Montiel, A. (2008). Identification of molecular determinants of channel gating in the transient receptor potential box of vanilloid receptor I. Faseb J 22, 3298–3309.

Vaught, A. (1996). Graphing with Gnuplot and Xmgr. Linux Journal.

Yao, J., and Qin, F. (2009). Interaction with phosphoinositides confers adaptation onto the TRPV1 pain receptor. PLoS Biol 7, e46.

